# Wild jackdaws learn to tolerate juveniles to exploit new information

**DOI:** 10.1101/2024.08.29.609864

**Authors:** Josh J. Arbon, Noa Truskanov, Emily Stott, Guillam E. McIvor, Alex Thornton

## Abstract

Social tolerance can enhance access to resources and is thought to be crucial in facilitating the evolution of cooperation, social cognition and culture, but it is unknown whether animals can optimise their social tolerance through learning. We presented wild jackdaws (*Corvus monedula*) with a novel social information problem using automated feeders: to access food, adults had to inhibit their tendency to displace juveniles and instead show tolerance by occupying an adjacent perch. Adults learned to tolerate juveniles, generalising across juveniles as a cohort, demonstrating learning of a new information-use strategy. The ability to learn to tolerate those that provide valuable information, and generalise across cohorts of informed individuals, may facilitate adaptive responses in the face of environmental change and help to explain the success of jackdaws in human-dominated environments.

## Introduction

Social tolerance, broadly defined as close proximity to conspecifics with limited aggression [1], is thought to be crucial to the evolution of prosociality [2] and social learning [3] in non-human animals, and underpins distinctive forms of cognition, cooperation and culture in humans [4]. Tolerating others can provide opportunities to learn important information such as how to access food [3], and has even been shown to help buffer the costs of ecological disasters [5], but social tolerance can also entail costs such as foraging competition [6]. General heuristics can aid in responding adaptively to environmental change[5], such as preferentially tolerating kin when resource availability is low [7]. However, if potential partners vary in the benefits they provide, then learning to tolerate specific individuals, or classes of individual, could be vital in facilitating access to information [8] or resources [9]. Despite these important implications, studies have yet to explicitly test whether animals in natural social networks can learn to tolerate others based on their value as sources of information.

Biases in social information use and social learning are well described across taxa [10]. For instance, many animals preferentially learn from older, more experienced individuals [11,12] that are more likely to provide reliable information. However, sources of useful information may change: in humans, for instance, younger generations commonly learn from their elders, but the digital revolution has flipped this pattern with older people seeking vital skills from younger “digital natives” [13]. Similar patterns could occur in animals. Juveniles are often easily displaced by adults and generally not used as preferential information sources [14,15], but juveniles are often more exploratory and innovative [16–19]. Adults may therefore benefit from learning to attend to and tolerate juveniles, especially in changeable or heterogenous environments where reliable models can change quickly [20–22]. Whilst some recent work indicates that animals adjust their social associations based on information about specific partners [23–25], whether these associations can be generalised to classes of individuals that share characteristics (e.g. juveniles) remains unknown. This combination of learning and generalisation that guide who to tolerate would represent the learning of a social information-use strategy [26]. We examined this possibility using an automated field experiment on wild jackdaws.

Jackdaws are social corvids that forage in fission-fusion social groups [27] and vary in their social tolerance of different partners, generally showing particularly high tolerance to close associates and kin [28]. They are proficient social learners about novel foods [29] and anthropogenic dangers [30], and have been shown to generalise learnt information in foraging contexts [15]. These abilities may contribute to their success in human-dominated habitats [22]. Previous work [23] shows jackdaws can learn to associate with individuals based on payoffs, but whether they can alter their tolerance or generalise this learning across individuals remains unknown. To answer these outstanding questions, we presented adults with a novel social information-use problem. Under normal conditions foraging adults do not preferentially associate with juveniles other than their own offspring (Supplementary Material: Age-class Assortment), nor attend to juveniles as sources of information [15]. We simulated a scenario where juveniles became ‘knowledgeable’ about a new foraging resource by allowing them free access to automated experimental feeders containing high-quality food (mealworms). As it took up to a couple of seconds for feeders to close, adults could often scrounge a small reward (max. one beakful) by aggressively displacing a juvenile as the feeder door closed. However, if adults refrained from displacing juveniles and instead tolerated their presence on the adjacent feeder, they gained the far larger reward of feeding *ad libitum* for the duration that they occupied the adjacent feeder (termed a cofeeding event: mean duration ± SE = 12.3 ± 1.5 s).

To test whether adults could learn to tolerate juveniles to exploit this novel scenario, we modelled changes in both cofeeding with juveniles and displacement of juveniles relative to adult experience. We included both the number of cofeeding events with a specific individual, as well as the number with all juveniles, to test between learning to tolerate specific individuals versus generalising learnt tolerance to juveniles more generally. We also placed pairs of feeders with low-quality food (grain), freely accessible to all birds, at distinct locations at the field-site, to test if changes observed at our experiment carried over into a related, but unrewarded, context.

## Methods

### Study System

Experimental sessions ran for five hours from ∼5:30 am between from 19 July to 13 August 2021, at our long-term jackdaw study population in Cornwall, UK (0°11’23”N, 5°10’54”W). The Cornish Jackdaw Project monitors a core nest-box population of ∼80 boxes as part of the wider jackdaw population, with more than 3000 individuals ringed over a 10-year period. Individuals are fitted with a unique set of coloured leg rings, one of which contains a Radio Frequency Identification (RFID) tag (IB Technologies, Leicester, UK), enabling remote detection via antennae at feeders. Approximately 150 chicks from nest boxes are ringed each year before fledging, and non-nestbox juveniles are ringed as part of routine trapping [31].

### Experimental Apparatus

Feeders automatically recorded the identity of any visiting RFID-tagged bird. Access to food was restricted by a motor-controlled door connected by a logger unit to an antenna in the perch (Naturecounters, Maidstone, UK; Figure 1a; Supplementary Material: Experimental Setup). Cofeeding events were determined from the temporal overlap between tag visits at adjacent, interconnected feeders (Figure 1a). Temporal data recorded which individual arrived first, initiating an event, and which individual subsequently joined. Displacements were characterised as events where a bird was detected arriving at a feeder within 2 seconds of a different bird departing (validated through video; see Supplementary Material: Displacement Validation).

**Figure 1.**
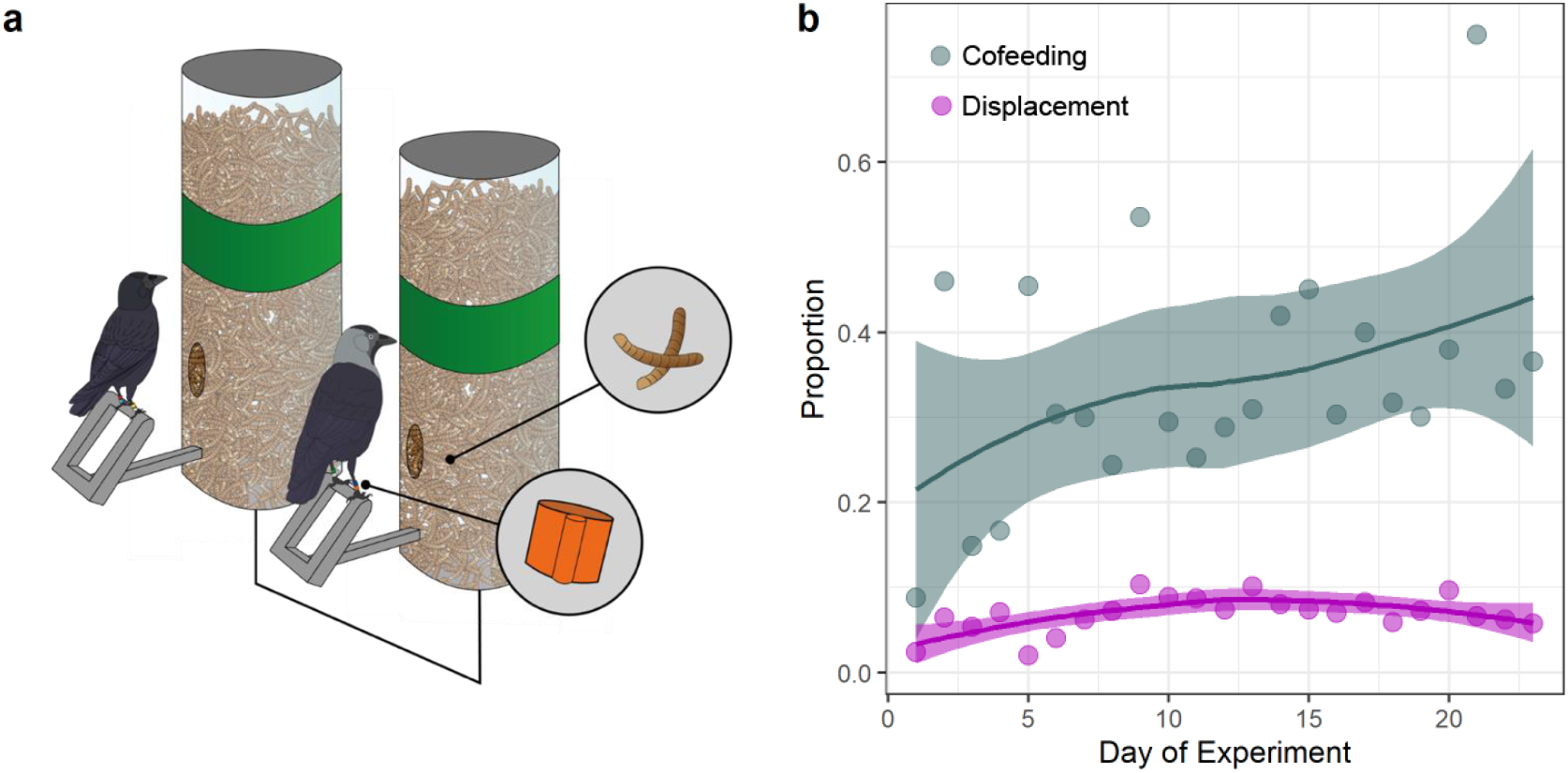
(a) Experimental feeders, showing an adult (right) in a successful cofeed with a juvenile. RFID tags (highlighted), determine access to mealworms by motor-controlled doors. (b) The daily proportion of visits where adults successfully co-fed with (grey-blue) or displaced (violet) a juvenile.

Feeders were split into two categories: *Experimental* feeders contained high-quality food (mealworms; known to be preferred by jackdaws [23]), whereas *passive* feeders contained low-quality food (grain). *Experimental* feeders were in connected pairs 0.5 m apart, with access to food determined by the tag identity and combination across both feeders (see Methods: *Treatments*). All tagged jackdaws could access *passive* feeders at all times. *Experimental* and *passive* feeders were visually distinguishable by visible food content and a green band present only on *experimental* feeders (Figure 1a).

We placed two pairs of *experimental* feeders at each of two sub-sites. These were flanked by a pair of *passive* feeders to create an experimental array. Together, the four feeders formed a shallow arc, each separated by 0.5 m (Supplementary Material: Experimental Setup). The *passive* feeders in experimental arrays were primarily designed to attract birds to the array, and to provide data on the individuals present in the area; data from these *passive* feeders within the experimental array are not presented in this study. For each experimental array, we also placed a corresponding pair of *passive* feeders in the same arrangement (0.5 m apart) elsewhere within the local area (∼50m away). These *passive* feeders were used to detect associations and displacements outside of the direct context of the experimental setup. Unfortunately, during the onset of the experiment, a dead jackdaw appeared next to one of the experimental arrays, resulting in a near-complete abandonment of all feeders at one sub-site (Supplementary Material: Feeder Visitation). As such, only data from feeders at one sub-site was analysed (N = 2 experimental arrays, 2 passive feeder pairs).

### Treatments

All RFID-tagged juveniles formed the ‘knowledgeable’ group, with unrestricted access to *experimental* feeders. Treatment adults (70% of adults chosen randomly) could access mealworm rewards from *experimental* feeders, but only if a juvenile was simultaneously present on the adjacent *experimental* feeder. Control adults (remaining 30% of adults) were not able to access food from the *experimental* feeders under any circumstances. All three groups had unrestricted access to *passive* feeders, including those within experimental arrays and the *passive* feeder pairs at nearby locations. The control adults were intended to provide baseline data on adult–juvenile interactions but visited experimental arrays at rates too low to allow meaningful analyses (Supplementary Material: Feeder Visitation). However, the use of a control group is not necessary for our key aim of detecting changes in within-individual behaviour over time (see Statistical Methods).

### Ethical Note

Research was conducted under ethical permission from the Penryn Campus Ethics Committee (eCORN000406), following Association for the Study of Animal Behaviour ethical guidelines [32]. Birds were ringed under BTO permits C6449, C5752, C6923 and C6079.

### Statistical Methods

We analysed data using Relational Event Models (REMs), a form of time-to-event model that accounts for the structure of the social network, to compare changes in the rates of cofeeding and displacement events based on individual experience (following Kings *et al*. 2023 [23]). REMs account for the order of events in a network, preventing data aggregation [33–35], and test hypotheses by comparing observed events against a set of ‘non-events’, permuted events that could hypothetically happen between individuals in the network; 10000 permutations were conducted ([36], see Supplementary Material: Relational Event Modelling for details).

We used this approach to separately model whether adults changed their propensity to (a) cofeed with juveniles (Supplementary Tables 1,3,5), and (b) displace juveniles (Supplementary Tables 2,4,6), based on their previous experience in the experiment. When investigating cofeeding likelihood, we considered the number of times an adult had previously cofed with a *specific individual* (*IRR individual*), as well as the number of times an adult had cofed with *any juvenile* (*IRR all juvs*) to investigate generalisation of learning. The likelihood of displacement was modelled as for cofeeding, with the addition of a fixed factor for the number of times an adult had displaced a juvenile (*IRR displacement*) to investigate whether learning from displacements reinforced this behaviour. By interacting these factors with the age (juvenile/adult) of the individual joined/displaced, we determined the likelihood of cofeeding with/displacing a juvenile relative to that of an adult as a function of experience. Results from experimental feeders were qualitatively identical when considering adult experience as either (a) the number of successful cofeeds where the adult joined a juvenile at an experimental feeder, or (b) the total number of successful cofeeds the adult participated in, both joining or being joined by a juvenile (Supplementary Tables 1–2). We present the results from (a) in the main text unless specified.

All results are reported as Incidence Rate Ratios, such that at IRR of 1.05 for a predictor variable means that the event type in question in 5% more likely for every unit increase in that predictor. We took the median model coefficient of the 10000 permuted models as a point estimate for each parameter, with the 2.5 and 97.5 percentiles of each coefficient taken to generate 95% confidence intervals.

## Results

### i) Experimental Results

As adult jackdaws gained experience, they increased their tolerance of juveniles. Over the experiment, we detected 2431 cofeeding events, where both *experimental* feeders were occupied simultaneously. Of those, 766 were successful cofeeds between 200 unique adult– juvenile dyads, 174 of which involved adults joining juveniles. Overall, the proportion of adult cofeeding events with juveniles (i.e., successes) roughly doubled across the course of the experiment (Figure 1b, grey-blue line).

At the individual level, the likelihood an adult joined a juvenile at the feeders (as opposed to joining another adult) increased by 3.4% for each previous successful cofeed the adult had participated in with any juvenile (*IRR all juvs* = 1.034, *CI =* 1.021–1.049; Figure 2a). Displacements of juveniles also became slightly more common over the course of the experiment (Figure 1b, violet line) as adults learnt that when displacing juveniles they could scrounge a small reward before the feeder closed (*IRR displacement =* 1.004, *CI* = 1.002–1.006). However, the likelihood that an adult displaced a juvenile (relative to another adult) declined by 2.2% per successful cofeed with any juvenile (*IRR all juvs =* 0.978, *CI* = 0.970–0.987; Figure 2b).

**Figure 2.**
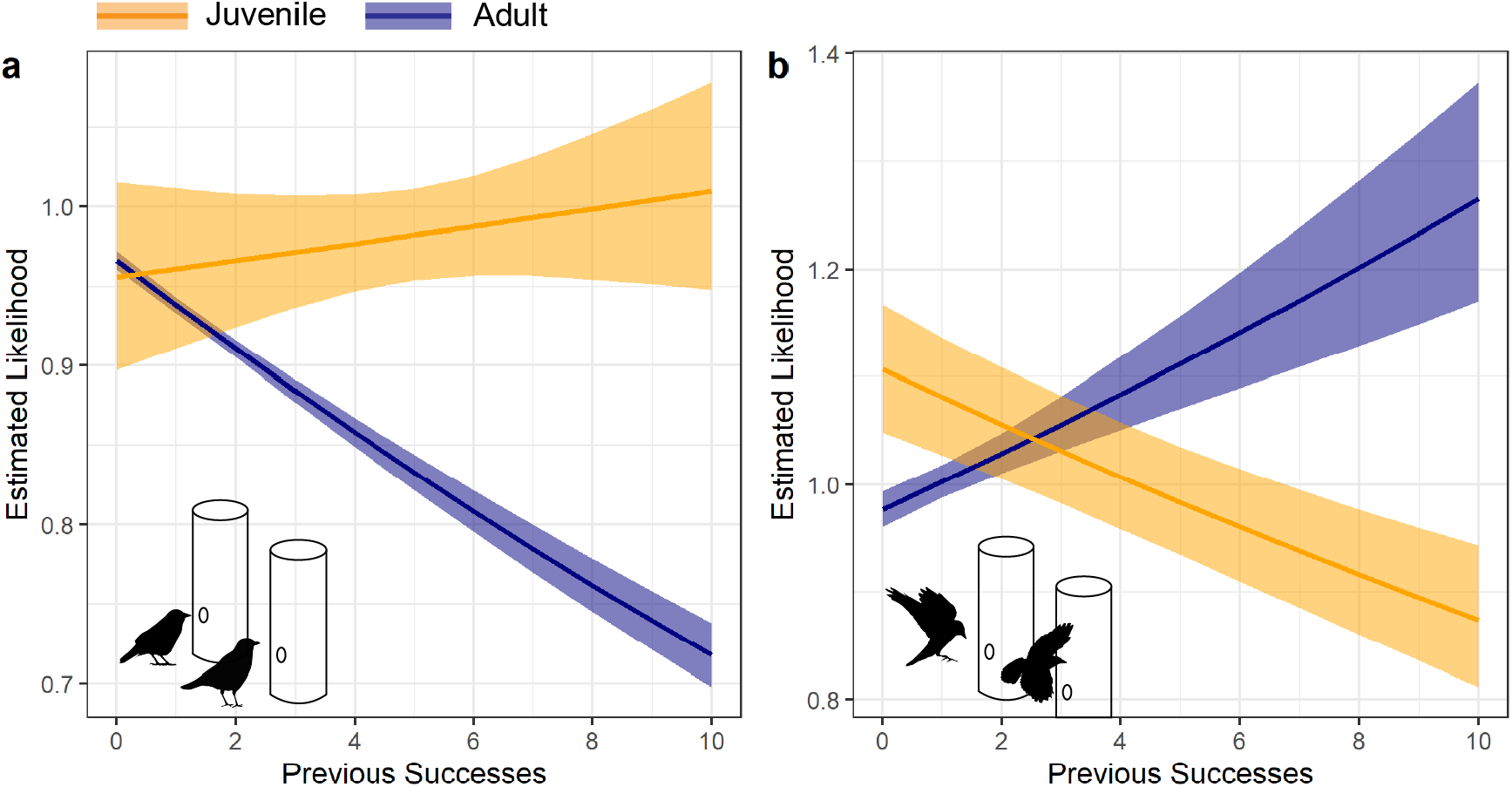
The difference in likelihood of an adult (a) cofeeding with, and (b) displacing a juvenile (orange) as opposed to another adult (blue) as adults acquired experience of successful cofeeding with all juveniles. Nine adults reached 6 or more previous successes. Lines and shaded area represent median and 95% REM estimates.

Together, these results indicate that successful cofeeding with any juvenile leads to adults learning to change their behaviour towards juveniles in general. Indeed, the effects of experience with specific juveniles (as opposed to any juvenile), are inconclusive on both the relative likelihood of cofeeding with (*IRR individual =* 1.002, *CI =* 0.986–1.020), or displacing (*IRR individual =* 0.992, *CI* = 0.983–1.001) a juvenile relative to another adult. Succeeding with a specific juvenile, or failing with a specific adult, appears to therefore be less important than overall successes with all juveniles.

One adult was responsible for a large number of successful joining events (37/174). If we exclude this adult from analyses, the effect of generalisation across juveniles becomes less clear for patterns of cofeeding; the estimate for the effect of experience with *all juveniles* is still positive, but the CI crosses 1 (i.e., no effect: *IRR all juvs* = 1.012, *CI* = 0.987–1.044), whilst the *specific individual* effect becomes more clear (*IRR individual* = 1.039, *CI* = 1.007–1.079). However, the effects for displacement, are qualitatively the same without the influential adult, with an increase in magnitude for the effect size of experience with all juveniles (*IRR individual* = 1.000, *CI* = 0.988–1.012, *IRR all juvs* = 0.952, *CI* = 0.938–0.968).

### ii) Carry-over of experimental manipulation

We also found limited evidence that tolerance learnt at the experimental feeders carried over to the non-experimental context. We did not see clear changes in cofeeding at passive feeders (Supplementary Table 3), however, adults were less likely to displace juveniles if their successes with all juveniles at the experimental feeders were higher (though only in models where successes included those where a juvenile joined an adult Supplementary Table 4).

### iii) Ruling out confounds

In principle, the apparent changes in adult behaviour with experience could arise as an artefact if adults with a greater pre-existing propensity to either associate with juveniles, or to visit experimental feeders, engaged with the experiment more and were more successful. We can rule this out, however as (1) adults’ preferences for cofeeding with juveniles at the start of the experiment were unrelated to the final number of successes they had and (2) adults that visited the experimental apparatus more frequently were not more successful overall (Supplementary Information: Initial Propensity).

## Discussion

Taken together, our results show that adult jackdaws can learn to tolerate juveniles when it is advantageous to do so. As they gained experience in the experiment, adults increased cofeeding with and reduced displacement of juveniles relative to adults, indicating they learned to tolerate juveniles based on their information value. Although there were small rewards available for displacing juveniles, these were largely outweighed by the benefits of tolerant cofeeding. Moreover, our results suggest generalisation across individuals may facilitate rapid, beneficial changes in tolerant behaviour, representing the learning of a new social information-use strategy, namely ‘tolerate juveniles’.

The potential benefits of such learned tolerance are abundant. Tolerating conspecifics has been shown to facilitate cooperation in ravens *Corvus corax* [2], whilst tolerance is implicated in the social learning of key foraging skills such as tool use in other corvid species [37,38], even in species not known to use tools in the wild [39]. Whilst information can be gathered from others without tolerance (e.g. broad-scale patch discovery by observing others from a distance), social tolerance can increase the chances of co-occurring with valuable individuals and observing their actions in close proximity. This may be especially valuable in species without fixed social groups, such as jackdaws and other animals living in fission-fusion societies, where individuals can switch to other social partners if they provide less aggression or better payoffs [23,27]. Our study therefore highlights the ability of jackdaws to flexibly alter information-use strategies through learning and generalisation. Whether the changes we detected in the context of foraging behaviour might carry over into other contexts (e.g. predator defence) is a major outstanding question. Addressing this will be challenging but important to help us understand how flexibility may help animals like jackdaws thrive in changeable, human-disturbed environments [21].

The findings that experience interacting with juveniles in general, over experience with specific individuals, was an important driver of behavioural change suggests that adults were able to generalise their learning to juveniles as a cohort. Whilst information-use strategies are known to be influenced by developmental stress [40], our findings show that learning can allow the rapid adjustment of strategies to novel conditions. This is consistent with suggestions that rather than evolving as fixed, species-typical traits [10], social learning strategies use domain-general associative learning processes to modify associations and inhibition of aggressive behaviour in response to reinforcement, enabling flexibility in the face of changing payoffs [26]. This flexibility also illustrates the benefits of tracking and responding to others’ behaviour in dynamic social environments, a key assumption of the Social Intelligence Hypothesis [41]. However, generalising among similar individuals could facilitate access to valuable new opportunities and information without needing to remember the outcomes of past interactions with many different individuals, reducing cognitive burdens.

It is also possible that different adults implemented different learning strategies. The nuanced changes in interpretation when removing the individual responsible for the largest number of successful joining events – i.e., strong evidence for generalising across juveniles when considering displacements but evidence for the importance of experience with specific individuals when considering co-feeding – highlights the potential for variation in who is attended to, and how. In practice it is likely that both remembering past interactions with specific individuals and generalising across groups of individuals will be used together to guide behaviour. Differences in how individuals integrate these processes could be underlaid by differences in cognitive abilities or prior experience [42], but we do not have a sample size large enough to investigate these differences robustly. Regardless of the specific strategy implemented by individuals, the key finding that adults learn to modify their social tolerance is clear: results cannot be explained by adults’ pre-existing preference for associating with juveniles, or their propensity to engage with the experiment in general.

Broadly, flexibly adjusting social tolerance, including through learning new information-use strategies, could facilitate the flow of information through networks by permitting close observation of a wide range of innovators. Indeed, the ability to capitalise on opportunities generated by innovators through flexible information-use is thought to be a crucial element of human culture [26]. Assessing the degree of flexibility in social information use and social learning by non-human animals will be an important factor in understanding how they respond to changes in the environment and increasing interactions with humans [43]. Our work also provides a rare illustration of the bi-directional interplay between information use and social structure in natural populations, contributing to our understanding of how sociality and decision-making processes coevolve [44,45].

## Acknowledgements

We would like to thank the Gluyas family and Odette Eddy for access to their land to conduct the experiment. Thanks to Andy Young for many valuable discussions and Stephanie King for comments.

## Author Contributions

Conceptualization: J.J.A. and A.T. Methodology: J.J.A. and A.T. Validation:

J.J.A. Formal Analysis: J.J.A. Investigation: J.J.A, N.T., E.S. Resources: J.J.A., G.E.M., and A.T. Data Curation: J.J.A. Writing – Original Draft: J.J.A and A.T. Writing – Review & Editing: J.J.A., N.T, E.S., G.E.M., and A.T. Visualization: J.J.A. Supervision: A.T. Project Administration: J.J.A., G.E.M., and A.T. Funding Acquisition: J.J.A. and A.T.

## Data accessibility

All data and code for this work can be found at https://figshare.com/s/34ce199d3d06a0be20b1.

