## Supplementary Figures and Tables for "Wild jackdaws learn to tolerate juveniles to exploit new information"

### Supplementary Material and Tables

#### Experimental Setup

Feeders were placed on wooden posts at a heigh of 1m from the ground, with the exception of one pair of *passive* feeders that were attached to the side of two adjacent tree trunks at a height of 2m to keep them out of the reach of cows in the adjacent field. Experimental arrays comprised a central pair of *experimental* feeders, flanked by a pair of *passive* feeders.

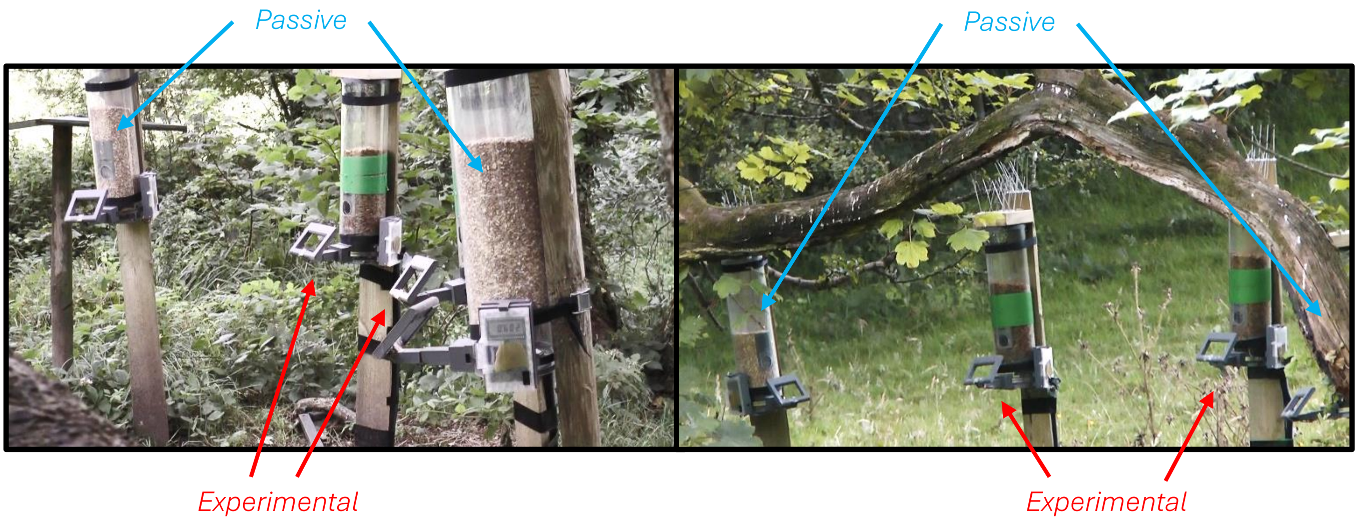


**Supplementary Figure 1.** Screengrabs from validation video cameras of the experimental arrays. Both arrays feature four feeders in a shallow arc, with two experimental, connected feeders flanked by a pair of passive feeders.

### Age-class Assortment

RFID feeder data from 2020 show that adults do not preferentially associate with juveniles under regular feeding circumstances. Instead, individuals were assorted by functional ageclass. Adults were more likely to be found in social groups with other adults, with similar within-age-class assortment for subadults (birds born the previous calendar year) and juveniles (birds born that year; Supplementary Figure 1). Gaussian mixture models were fitted to RFID data to determine social groups [1], with dyadic association calculated via simple ratio index. Assortment was then calculated using the assortment.discrete function in the *assortnet* package [2]. Data-stream permutation was conducted to generate null models of assortment, and the observed age-class assortment was significantly higher than that generated from permuted datasets ($P < 0.05$, $N_{Rands}$ = 1000).

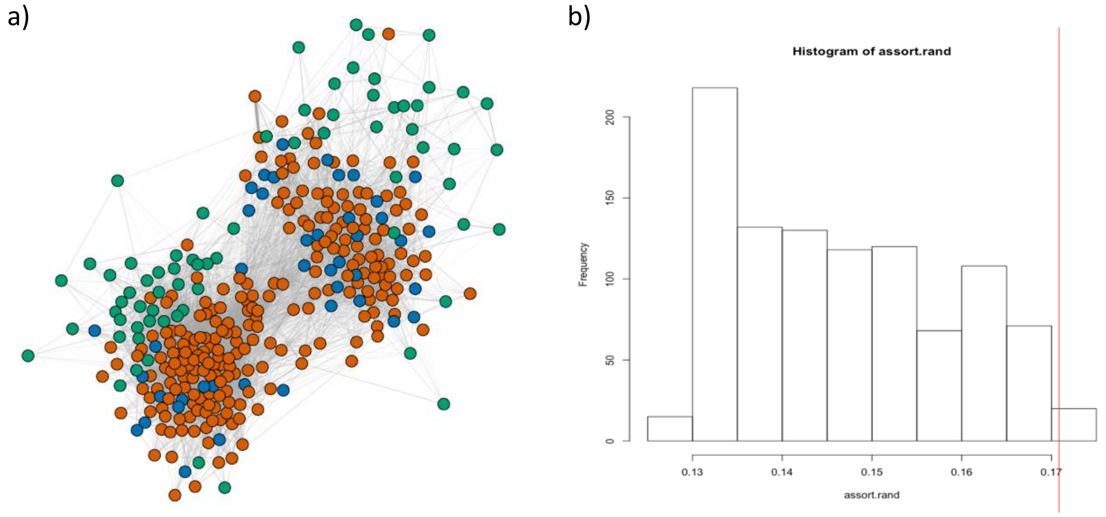


**Supplementary Figure 2.** (a) Visualisation of RFID feeder network from 2020, with node colours representing the three functional age-classes of jackdaws: orange = adult, blue = subadult, green = juvenile. Thickness of lines between individuals represents the strength of association between the individuals. (b) Histogram of permuted assortment by age-class values for the network, with red line representing observed assortment, illustrating the higher-than-expected assortment by age-class in the network.

### Feeder Visitation

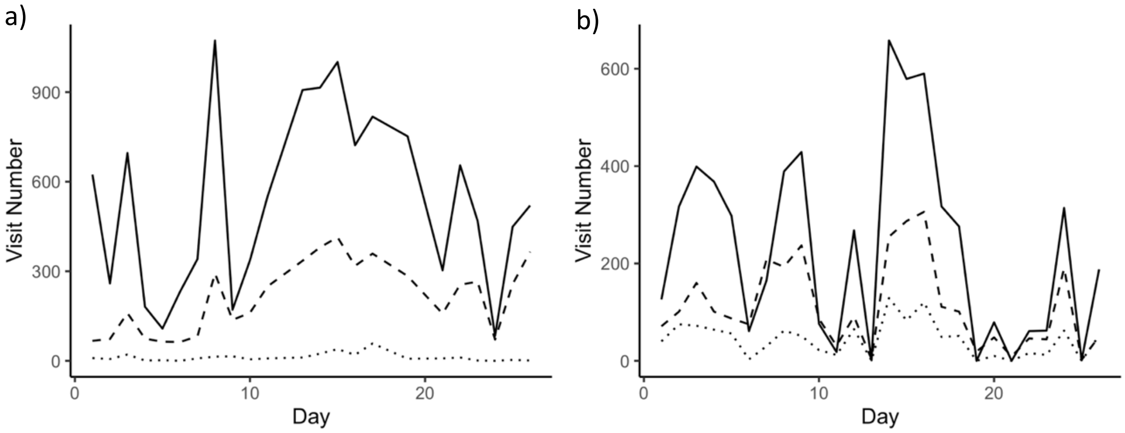


**Supplementary Figure 3.** Number of feeder visits by day at (a) experimental, and (b) passive feeders over the course of the experiment. Solid line represents treatment adults, dashed line juveniles, with dotted line control adults.

Supplementary Figure 2 displays feeder visitation at the sub-site that was carried forward for analysis. As highlighted in the main text, in the initial days of the experiment, a farmer discovered a dead, ringed, jackdaw at the other sub-site and placed it next to an experimental feeder array at that sub-site whilst it was in operation, so we would easily find the bird and could therefore record the death and remove the rings. This well-intentioned action appears to have inadvertently created an association between a deceased bird and feeders at the sub-site, as following this event all the feeders at this sub-site, both the array adjacent to the dead bird and those >100 m away, received next to no visits going forward. After 10 further days, the majority of which contained no visits by any birds, we ceased operating the feeders at this sub-site for the duration of the experiment.

### Initial Propensity

As illustrated in Figure S3 below, the results presented in the main text are cannot be explained by adult individuals' propensity to associate with juveniles.

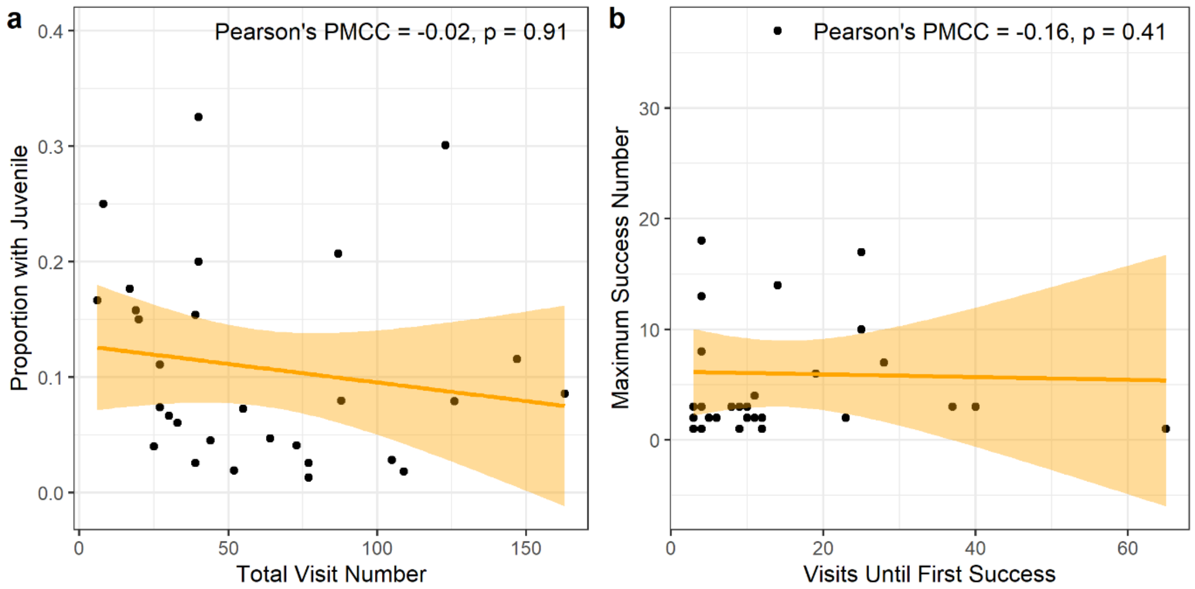


**Supplementary Figure 4.** (a) The total number of visits adults made to experimental feeders was not related to the total number of visits made with juveniles (successful cofeeds). This shows that 'juvenile-friendly' adults did not engage with the experiment more often and thus experimental results cannot be explained by pre-existing preferences for associating with juveniles. (b) The number of failures an individual had prior to its first success – a proxy for propensity to associate with a juvenile when the adult had no personal experience of experimental payoffs – was not related to the ultimate number of successful cofeeds an individual participated in. This provides further evidence that adults that were more likely to associate with juveniles to begin with were not driving the results by gaining more successes in total.

### Displacement Validation

To validate that changes of identity within a short time window were being correctly identified as displacements, we transcribed a 10-hour section of video of interactions at feeders (as shown in Supplementary Figure 1). By applying a variety of time windows, a threshold of two seconds was identified as the optimal trade-off between (i) ensuring a large number of displacements were included: as birds often had to adjust their posture having displaced another, the antennae did not always read the new tag immediately, and (ii) minimizing false positive detections: i.e. instances where a subordinate bird arrived immediately after a more dominant bird left (akin to scrounging) but no displacement occurred. It was especially important to avoid false positives because the signals for displacements and non-aggressive replacements would look identical in the RFID data stream, despite the underlying behavioural process being fundamentally different. At a threshold of two seconds, 50% of displacements were automatically detected, with a false positive rate of less than 5% (132 decoded from video, 65 correctly identified in data-stream, with 3 false positives). We therefore chose this trade-off between false negatives and false-positives to maximise the avoidance of false positives. Whereas false positives could bias results (by incorrectly classifying replacements of juveniles as displacements) false negatives (failures to detect displacements) are extremely unlikely to produce systematic biases in within-individual changes of behaviour over the course of the experiment.

### Relational Event Modelling

Non-events for REMs are conventionally evenly sampled from the pool of individuals [3–5], which is not a realistic assumption for data of interactions between freely interacting wild animals, where some individuals are more represented in the data-stream than others. In line with conventional social network approaches [6,7], we used a permutation method to generate null datasets that maintained the structure of the data regarding individual activity at the experiment, as well as the opportunities to succeed at the task based on the other individuals present, something we would be unable to incorporate into traditional modelling approaches. We created 10000 permuted datasets by randomising the identity of the source (individual joining a cofeed or displacing) for all events where this source individual was an adult; this method therefore crucially controls for the opportunity to join/displace. This results in a dataset where for each observed event there is a corresponding permuted 'non-event'; each event, both observed and permuted, has covariates that can be modelled against the event type (i.e. with an adult or with a juvenile). We then ran cox proportional hazard models using the survival package [8] in R v4.4.0 [9] comparing each permuted dataset (non-events) and the observed data, including a strata term for each day of the experiment to control for the baseline rate of observed events. This method enables us to ask questions such as: given the structure of visits and the opportunities to join juveniles and adults, was an adult that has experience cofeeding with juveniles more or less likely to join a juvenile compared to joining an adult, than an adult that had little experience cofeeding with juveniles? This approach also enables us to separate experience of the individual (experience cofeeding with any juvenile) from the characteristics of the dyad (number of times that specific dyad has cofed), to test specific questions relating to generalisation of learnt information.

### Model Output Tables

#### Model Term Definitions

**Target.juvenile**: Binary factor representing whether the individual cofed with or displaced was a juvenile (1) or another adult (0)

**Adult.successes**: Number of previous successful cofeeds with any juvenile prior to this event. Represented by '*IRR all juvs*' in main text.

**Dyad.cofeeds**: Number of cofeeding events between this specific dyad of individuals prior to this event. Represented by '*IRR individual* ' in main text.

**Juvenile.displacements**: The number of displacements of juveniles carried out prior to this event. Represented by '*IRR displacement* ' in main text.

Such that **Target.juvenile * Adult.successes** represents the estimated change in likelihood of the target being a juvenile for each increase in the number of previous successful cofeeds an adult has participated in. An IRR of 1.05 for this term therefore predicts a 5% increase in the likelihood of a juvenile being the target for every one increase in adult prior success. Similarly, **Target.juvenile * Juvenile.displacements** represents the change in likelihood of an adult displacing a juvenile for each prior displacement of a juvenile that adult has participated in. All interaction terms where the 95% bounds do not cross 'no effect' are highlighted in bold.

Models are analysed with **Adult.successes** calculated using two related metrics. Models presented in (a), and in the main text, are analysed using the the number of successful cofeeds the adult joined at an experimental feeder, whereas models in (b) use the total number of successful cofeeds the adult had participated with, including those in (a) and those where the adult was present first and then joined by a juvenile. Both joining a juvenile and being joined by a juvenile at the experimental feeders represent potential opportunities to learn the contingency between juveniles and food. Whereas waiting to be joined could be a strategy employed by adults, this gives them little agency to control who joins them. Analysing events where one of the two perches is already occupied before the adult makes a decision therefore provides a scenario where we can test adult performance.

**Supplementary Table 1.** Likelihood of an adult joining a cofeeding event at experimental feeders. Model output given for Adult.successes calculated as a) the number of successful cofeeds the adult joined a juvenile at an experimental feeder, and b) the total number of successful cofeeds the adult participated in, both joining or being joined by a juvenile.

| *a) Joined Successes* | *2.5% IRR* | *Median IRR* | *97.5% IRR* |
|---|---|---|---|
| Target.juvenile | 0.937 | 0.997 | 1.060 |
| Adult.successes | 0.962 | 0.970 | 0.973 |
| Dyad.cofeeds | 1.007 | 1.007 | 1.008 |
| **Target.juvenile * Adult.successes** | **1.021** | **1.034** | **1.049** |
| Target.juvenile * Dyad.cofeeds | 0.986 | 1.002 | 1.020 |
| *b) All Successes* | | | |
| Target.juvenile | 0.925 | 0.988 | 1.055 |
| Adult.successes | 0.995 | 0.993 | 0.996 |
| Dyad.cofeeds | 1.006 | 1.007 | 1.007 |
| **Target.juvenile * Adult.successes** | **1.005** | **1.008** | **1.012** |
| Target.juvenile * Dyad.cofeeds | 0.976 | 0.993 | 1.012 |

**Supplementary Table 2.** Likelihood of an adult performing a displacement at experimental feeders. Model output given for Adult.successes calculated as a) the number of successful cofeeds the adult joined a juvenile at an experimental feeder, and b) the total number of successful cofeeds the adult participated in, both joining or being joined by a juvenile.

| *a) Joined Successes* | *2.5% IRR* | *Median IRR* | *97.5% IRR* |
|---|---|---|---|
| Target.juvenile | 0.985 | 1.037 | 1.091 |
| Adult.successes | 1.005 | 1.010 | 1.016 |
| Dyad.cofeeds | 1.004 | 1.005 | 1.007 |
| Juvenile.displacements | 0.995 | 0.997 | 0.998 |
| **Target.juvenile * Adult.successes** | **0.970** | **0.978** | **0.987** |
| Target.juvenile * Dyad.cofeeds | 0.983 | 0.992 | 1.001 |
| **Target.juvenile * Juvenile.displacements** | **1.002** | **1.004** | **1.006** |
| *b) All Successes* | | | |
| Target.juvenile | 0.992 | 1.044 | 1.099 |
| Adult.successes | 1.003 | 1.004 | 1.006 |
| Dyad.cofeeds | 1.004 | 1.005 | 1.007 |
| Juvenile.displacements | 0.994 | 0.995 | 0.997 |
| Target.juvenile * Adult.successes | **0.988** | **0.991** | **0.994** |
| Target.juvenile * Dyad.cofeeds | 0.989 | 0.998 | 1.008 |
| Target.juvenile * Juvenile.displacements | **1.004** | **1.006** | **1.009** |

**Supplementary Table 3.** Likelihood of an adult joining a cofeeding event at passive feeders. Model output given for Adult.successes calculated as a) the number of successful cofeeds the adult joined a juvenile at an experimental feeder, and b) the total number of successful cofeeds the adult participated in, both joining or being joined by a juvenile.

| *a) Joined Successes* | *2.5% IRR* | *Median IRR* | *97.5% IRR* |
|---|---|---|---|
| Target.juvenile | 0.964 | 1.010 | 1.051 |
| Adult.successes | 0.985 | 0.995 | 1.009 |
| Dyad.cofeeds | 1.004 | 1.010 | 1.012 |
| Target.juvenile * Adult.successes | 0.909 | 1.018 | 1.212 |
| Target.juvenile * Dyad.cofeeds | 0.734 | 0.906 | 1.248 |
| *b) All Successes* | | | |
| Target.juvenile | 0.978 | 1.027 | 1.070 |
| Adult.successes | 0.999 | 1.000 | 1.003 |
| Dyad.cofeeds | 1.003 | 1.010 | 1.011 |
| Target.juvenile * Adult.successes | 0.962 | 0.988 | 1.031 |
| Target.juvenile * Dyad.cofeeds | 0.792 | 0.972 | 1.360 |

**Supplementary Table 4.** Likelihood of an adult performing a displacement at passive feeders. Model output given for Adult.successes calculated as a) the number of successful cofeeds the adult joined a juvenile at an experimental feeder, and b) the total number of successful cofeeds the adult participated in, both joining or being joined by a juvenile.

| | | | |
|---|---|---|---|
| *b) Joined Successes* | | | |
| Target.juvenile | 0.978 | 1.013 | 1.049 |
| Adult.successes | 0.998 | 1.016 | 1.038 |
| Dyad.cofeeds | 1.012 | 1.024 | 1.027 |
| Juvenile.displacements | 0.995 | 0.998 | 1.001 |
| Target.juvenile * Adult.successes | 0.816 | 0.919 | 1.040 |
| Target.juvenile * Dyad.cofeeds | 0.844 | 0.937 | 1.088 |
| Target.juvenile * Juvenile.displacements | 0.994 | 1.007 | 1.022 |
| *a) All Successes* | *2.5% IRR* | *Median IRR* | *97.5% IRR* |
| Target.juvenile | 0.983 | 1.019 | 1.056 |
| Adult.successes | 1.000 | 1.004 | 1.007 |
| Dyad.cofeeds | 1.012 | 1.024 | 1.027 |
| Juvenile.displacements | 0.995 | 0.998 | 1.001 |
| **Target.juvenile * Adult.successes** | **0.938** | **0.963** | **0.988** |
| Target.juvenile * Dyad.cofeeds | 0.878 | 0.980 | 1.139 |
| **Target.juvenile * Juvenile.displacements** | **1.000** | **1.013** | **1.027** |

***Supplementary Table 5.*** Experimental feeder cofeeding model (as in Supplementary Table 1a) with overrepresented individual removed.

| | *2.5% IRR* | *Median IRR* | *97.5% IRR* |
|---|---|---|---|
| Target.juvenile | 0.894 | 0.979 | 1.073 |
| Adult.successes | 0.934 | 0.953 | 0.958 |
| Dyad.cofeeds | 1.007 | 1.008 | 1.009 |
| Target.juvenile * Adult.successes | 0.987 | 1.012 | 1.044 |
| Target.juvenile * Dyad.cofeeds | **1.007** | **1.039** | **1.079** |

**Supplementary Table 6.** Experimental feeder displacement model (as in Supplementary Table 2a) with overrepresented individual removed.

| | *2.5% IRR* | *Median IRR* | *97.5% IRR* |
|---|---|---|---|
| Target.juvenile | 1.013 | 1.075 | 1.138 |
| Adult.successes | 1.017 | 1.025 | 1.034 |
| Dyad.cofeeds | 1.004 | 1.006 | 1.009 |
| Juvenile.displacements | 0.994 | 0.996 | 0.997 |
| **Target.juvenile * Adult.successes** | **0.938** | **0.952** | **0.968** |
| Target.juvenile * Dyad.cofeeds | 0.988 | 1.000 | 1.012 |
| **Target.juvenile * Juvenile.displacements** | **1.003** | **1.005** | **1.008** |
